# SYNAPTIC INTERACTIONS IN PRIMATE MOTOR CORTEX: RELATIONS BETWEEN CONNECTIVITY AND INTRACORTICAL LOCATION

**DOI:** 10.1101/2025.04.22.650058

**Authors:** Michikazu Matsumura, Dao-fen Chen, Eberhard E. Fetz

**Affiliations:** Department of Neurobiology and Biophysics, and Regional Primate Research Center, University of Washington, Seattle WA 98195 USA

**Keywords:** primate motor cortex, intracellular recording, spike-triggered averaging, synaptic connectivity, cortical circuit

## Abstract

To investigate intracortical microcircuits within primate motor cortical areas, we documented pairs of neurons whose synaptic interactions were identified by spike-triggered averaging (STA) of intracellularly recorded membrane potentials and whose relative location was histologically identified.

Average synchronous excitation potentials (ASEPs) were the most commonly observed feature in STAs between neuron pairs in all cortical layers (about 70%). This synchrony spread broadly for distances of more than 4 mm, gradually decreasing in amplitude and probability with cell separation.

Excitatory postsynaptic potentials (EPSPs) were observed among 9% of the neuron pairs, whose separation extended for distances of more than 4 mm. The peak amplitudes of EPSPs were inversely correlated with cell separation. Most source neurons were located in layer II-III or layer V, while the postsynaptic target neurons had wide laminar distribution.

The probability of finding excitatory connections was not uniform within the cortical space. The connectivity between neuron pairs was relatively dense within 1.0 mm, and became sparse for distances between 1.0 and 2.0 mm, and showed a second peak for distance between 2.0 and 4.0 mm.

Inhibitory postsynaptic potentials (IPSPs) and/or average synchronous inhibitory potentials (ASIPs) were observed for 10% of neuron pairs. Most of these were separated by less than 1.0 mm, suggesting that their influence was restricted within their own and neighboring columns. The major source neurons that provide these inhibitory effects were located within layer II-III. We conclude that excitatory synaptic effects (ASEPs and EPSPs) are widely distributed and omni-directional within primate motor cortex, while inhibitory connections (ASIPs and IPSPs) are restricted within columnar dimensions, and are predominantly directed from superficial to deeper layers.

The appearance of independent zones that had no functional connectivity with neighboring columns indicates that primate motor cortical areas may not be organized in a uniform way. This would support the sparse coding theory and multiple representations of cortical output in the motor cortex.

## INTRODUCTION

Recent anatomical studies have revealed the structural substrate for synaptic interactions within cerebral cortex (Aroniadou & Keller 1993, Binzegger et al 2004, Lund et al 1993, Peters & Jones 1984, White & Keller 1989). Afferent fibers to cortex spread over 4 mm horizontally (Strick & Sterling 1974), suggesting that cells in cerebral cortex receive common synaptic inputs. Intracortical spread of axon branches has been revealed by degeneration studies (Shanks et al 1978) and through injection of horseradish peroxidase (HRP) into the cortex (DeFelipe et al 1986, Gilbert & Wiesel 1983, Huntley & Jones 1991). Axon collaterals of pyramidal cells tend to spread and terminate in discrete columns, suggesting that excitatory synaptic potentials also affect broader but discrete columns. Most of these anatomical studies found a non-uniform distribution of axonal branching. Histochemical studies show that cortical interneurons are predominantly GABAergic, and have relatively short axonal distribution (Fish et al 2018). GABAergic interneurons called ‘chandelier cell’ located within layer II and III have axon terminals directly on initial segment and axon hillock of layer V pyramidal cells (Wang et al 2016).

These histological studies indicate the extent over which synaptic interaction could occur within the cortex. However, morphological studies cannot provide any quantitative information about the nature and strength of physiological interactions. Synaptic interactions between cortical neurons can be documented by cross-correlation techniques (Fetz et al 1991, Murphy et al 1985, Ts’o et al 1986). Both anatomical and physiological information of the same cells is needed to better elucidate intracortical circuitry. Such combined information has been obtained through appropriate in vitro studies (Thomson & Lamy 2007).

We investigated both anatomical locations and functional synaptic interactions of cells in primate motor cortical areas. Clarifying these functional relationships is essential to understand the neural mechanisms that mediate intracortical computations. We previously described the kinds of synaptic interaction between neurons in primate motor cortical areas: spike triggered averages (STA) of membrane potentials from simultaneously recorded neuron pairs revealed four basic features: excitatory and inhibitory postsynaptic potentials (EPSPs & IPSPs), and synchronous excitatory and inhibitory potentials (ASEPs & ASIPs) (Matsumura et al 1996). The present study adds histological information on the locations of the neurons showing these functional synaptic interactions. In addition, we also include neuron pairs whose STAs revealed no synaptic features, to assess the relative probability of these functional connections within the cortex.

## METHODS

The present experiments were approved by the University of Washington Animal Care Committee and performed according to NIH guidelines for animal use.

Recording procedures and data analysis were described previously (Matsumura et al 1996) and the present data comprise the part of that database for which electrophysiological results and confirmed recording sites were both available. The present data were obtained from 5 hemispheres of 4 macaque monkeys. Para-sagittal sections of the recording areas were made to reconstruct the recording sites.

### Surgical preparation and recording

For the initial sterile implant surgery, the monkeys were deeply anesthetized with sodium pentobarbital (Nembutal, Abbott, 35 mg/kg, i.p.) or with Halothane. The skull was exposed and 10-12 small vitallium screws were implanted. Two stainless steel tubes were cemented in parallel on the skull over frontal and occipital areas and subsequently used to attach the head to the primate chair. A concentric bipolar electrode was implanted for stimulating the pyramidal tract at the medullary level (AP = 0.0 mm, L = 1.5 mm). The exposed skull was covered with a thin layer of dental acrylic and the monkey was treated with antibiotics to prevent infection.

Recording sessions began after at least 7 post-operative days of recovery. On each recording day, the monkey was given a small dose of Ketamine (0.5 mg/kg, i.m.) and anesthetized with Halothane (1%, with 2-3 l/min oxygen and 1 l/min nitrous oxide). The monkey was seated in the recording chair, and the head attached to a stereotaxic frame via the implanted tubes (for details, see Fig. 1 of Matsumura 1979). The stereotaxic frame provided support and repeatable stereotaxic coordinates for electrode carriers. A small elliptical hole (about 1.5 × 2.5 mm) was drilled through the acrylic and the skull at site within the region bounded by A10 to A20 and L5 to L20 (this area covers the anterior portion of the central gyrus). Subsequent histology confirmed that these sites included area 4 and area 6. The dura was incised with a 26 g. hypodermic needle to expose the surface of the cortex.

**Figure 1.**
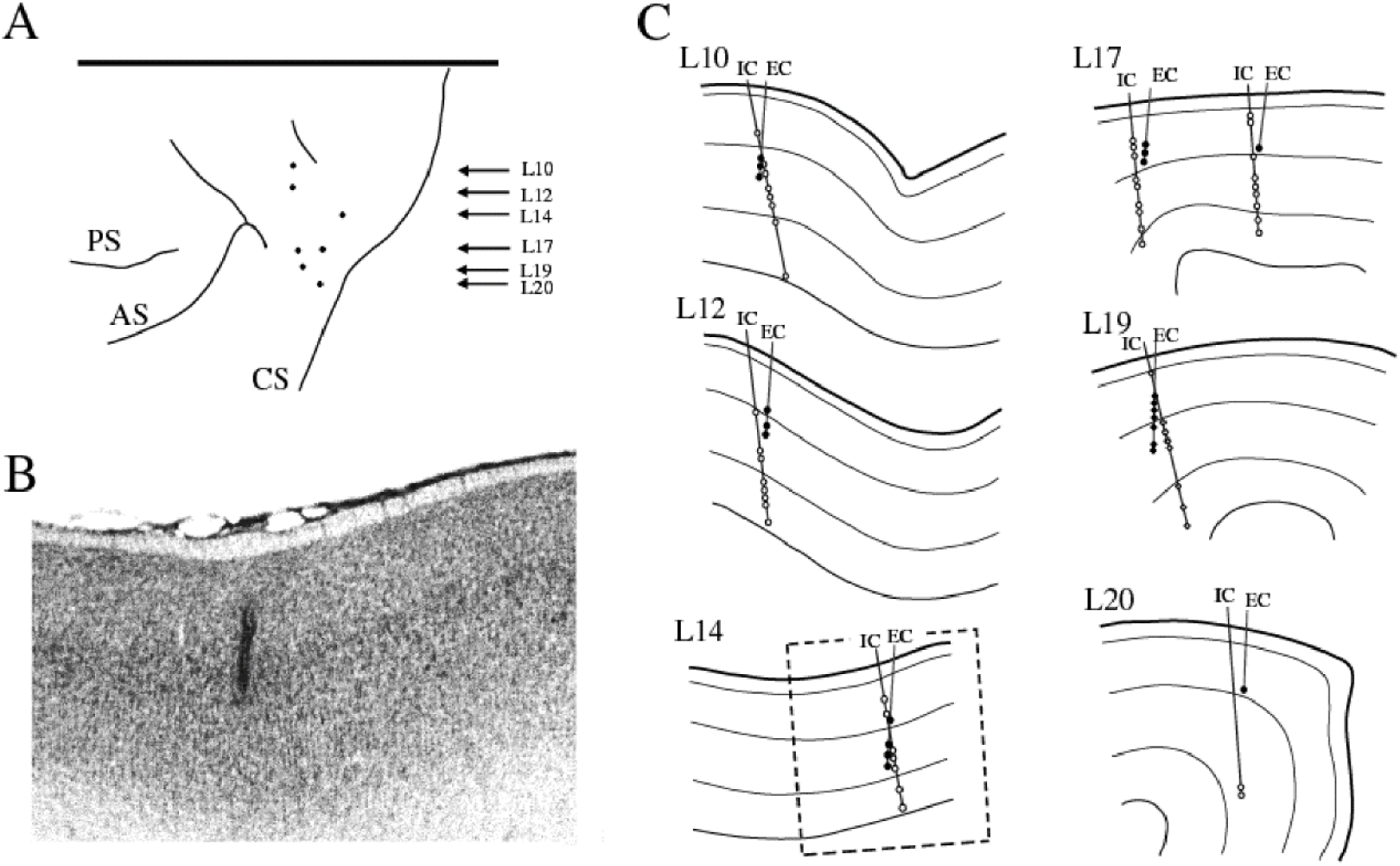
Histological reconstruction of sites of the recorded neuron pairs. A: surface view of the recording sites obtained from one hemisphere, from which parasagittal sections were analyzed. Arrows show the laterality of the recordings in mm. B: histological record of the recording site at L14. C: reconstruction of serial sections containing electrode tracks, obtained at identified laterality. Sites where intracellular (IC) and extracellular (EC) recording were obtained are indicated by open and filled circles, respectively. Roman numerals indicate the cortical layer. The broken-line rectangle in L14 marks the region shown in B.

Electrodes for intracellular (IC) and extracellular (EC) recording were inserted into the cortex with independently movable stereotaxic carriers (David Kopf) with the aid of a binocular microscope. The EC electrode was inserted in a vertical stereotaxic direction, and the IC electrode was inserted at an angle of 10 – 15° from vertical in the parasagittal plane. The IC electrode was advanced by a pulse-stepping microdrive (Burleigh Inchworm). To dampen cortical pulsation the hole in the skull was filled with 4% agar dissolved in saline after electrode tips were placed in the superficial cortical layer, and the monkey was allowed to recover.

Intracellular recordings were obtained with glass micropipettes (2 mm outer diameter) filled with K-Methylsulfate, with resistance between 10 and 40 Megohms. The EC electrode was a multi-barreled glass micropipette. One barrel of the pipettes contained a carbon fiber of 7 mm diameter (Toray, Toreca-3000) (Armstrong-James and Millar 1979). Five to fifty microns of the carbon fiber was exposed at the tip of the pipette, and electric signals were led to an amplifier via NaCl medium. The other barrel was filled with Na-glutamate (10 mM) to activate the isolated single unit(s) by micro-iontophoretic application of glutamate.

The carbon fiber barrel was connected to an AC amplifier, and the iontophoretic side to a constant-current isolation amplifier for application of glutamate by anodal current of up to 70 nA (Axon Instrument, Axoprobe). A small platinum-plated pin implanted near the recording site was used as a reference electrode. The electric signals from the IC electrode were led to a high-gain monitor. All signals were recorded on a 14-channel FM tape recorder (Honeywell 100).

### Spike-triggered averaging

Simultaneous recordings of intracellular (IC) membrane potential and extracellular (EC) unit activity from pairs of cortical neurons were recorded on analog magnetic tape for off-line analysis by STA. In many cases, particularly in anesthetized monkeys, the EC neurons were activated by iontophoretically applied Na-glutamate. Iontophoretic activation would favor somatic or dendritic recording and exclude the possibility of evoking activity from passing fibers.

A computer compiled spike-triggered averages of the IC membrane potential and included events preceding the trigger. Time-amplitude window discriminators (BAK, DIS-1) generated pulses from action potentials in the EC and IC records. The EC pulses were used to trigger the STA. The IC pulses were used to reject sweeps in the STA that contained IC action potentials and were also used to compile cross-correlograms of EC and IC spikes. The minimum number of sweeps to detect monosynaptic connection in a STA was taken as 100, although most averages included many more events. In a few exceptional cases, the STA revealed sufficiently large potentials to judge synaptic connections with less than 100 sweeps. IC recordings from neurons with large baseline fluctuations due to cardiac or respiratory pulsations were excluded from analysis.

### Histology and reconstruction of the recording sites

To mark recording sites, anodal and cathodal currents of 10 µA were passed for 10 sec through the carbon fiber to make coagulate deposits at EC sites (Sawaguchi et al 1986). After each recording session (lasting 2 to 6 hours), the electrodes were withdrawn from the cortex without changing the electrode carrier positions. With the monkey lightly anesthetized with Ketamine, antibiotics were applied to the cortex and the skull opening was closed with dental cement. The monkey was returned to its cage, and allowed at least one day for recovery between recording sessions.

The three-dimensional configuration of the two electrodes was measured after each recording by repositioning the electrodes at the coordinates of the recordings. The angles of the electrodes and distances between their tips at the closest crossing point were measured with the aid of binocular magnification at every recording position, and these coordinates were later compared with histological sections.

Direct distances between the pair recording sites were determined by trigonometric calculation based on the depth readings of the micro-drive from the cortical surface, and the angles between the electrodes, as well as the information of the crossing point of the two electrodes. Since many pairs of penetrations were not placed exactly within a same parasagittal section, the separation of the parasagittal plane was also taken into account as the z-axis value.

The distance measurements became less accurate when the distance in the z-axis was wider or the recording sites were located further laterally, since the parasagittal plane would deviate from columnar vertical orientation. In these lateral sites, the exact laminar location could not be determined as accurately. In those cases, only the measurement of direct distance between the EC-IC sites was accepted for the database. However, if the laminar location of the recording sites was in doubt, the data were excluded from laminar analysis.

Horizontal distances between the EC-IC sites along the columnar structure were calculated by taking account of the cortical geometry. The horizontal distance measurement was normalized by projecting coordinates onto the middle of layer V. Five hundred µm was regarded as a single columnar distance, and used as a unit distance in the following description.

At the end of the experiment, the monkeys were deeply anesthetized with Nembutal and perfused by saline followed by 10% formalin. The brain was post-fixed in 30% sucrose-formalin solution. Before the brain tissue was cut for histology, the surface was photographed and used for subsequent reconstruction. The tissue was cut either coronally or parasagitally into 200 μm serial sections, and stained by standard Nissl methods. The entry points of the recording electrodes were easily detected by connective tissue on the cortical surface and concentrated aggregations of glia in layer I and remaining pia mater. The electrode tracks were recognizable by glia or blood cells along the penetrating electrode track. The recording sites were identified with the aid of the depth reading of the electrode carrier and carbon deposits in the section.

## RESULTS

### Recording sites and their laminar locations

In 5 hemispheres of 4 monkeys we recorded from 220 pairs of EC and IC neurons within the motor and premotor cortex for sufficiently long durations to assess features of synaptic interactions in STAs. Of these 220 pairs, recorded in 22 pairs of penetration tracks, 182 EC-IC pairs recorded in 18 penetrations were histologically recovered and their laminar location identified.

Figure 1A shows a surface view of the recording sites obtained from one hemisphere, which was sectioned parasagittally. Fig. 1B shows an example of the histology of the recording site obtained at laterality L14. Layer V is clearly distinguishable by the presence of numerous giant pyramidal cells. The vertical thick band, from layer III to V, indicates hematological signs of the EC electrode penetration.

The electrode tracks obtained from each laterality are superimposed on the serial sections in Fig.1C. Open and filled circles indicate sites where intracellular (IC) and extracellular (EC) recordings were obtained, respectively. In all penetration tracks the numbers of EC sites are less than those of IC sites because many IC recordings were obtained with stationary positioned EC recording electrodes. Thin lines indicate the border of the layers. The border between layers II and III was barely distinguishable, so these were combined unless otherwise noted.

### Types of synaptic interaction and their laminar location

As previously described (Matsumura et al 1996), STA revealed four basic types of synaptic interaction, which could also occur in combination. Fig. 2 compares the average shapes of the 4 types of spike-triggered effects, drawn according to the mean latencies and amplitudes given in Table 2 of (Matsumura et al 1996).

**Figure 2.**
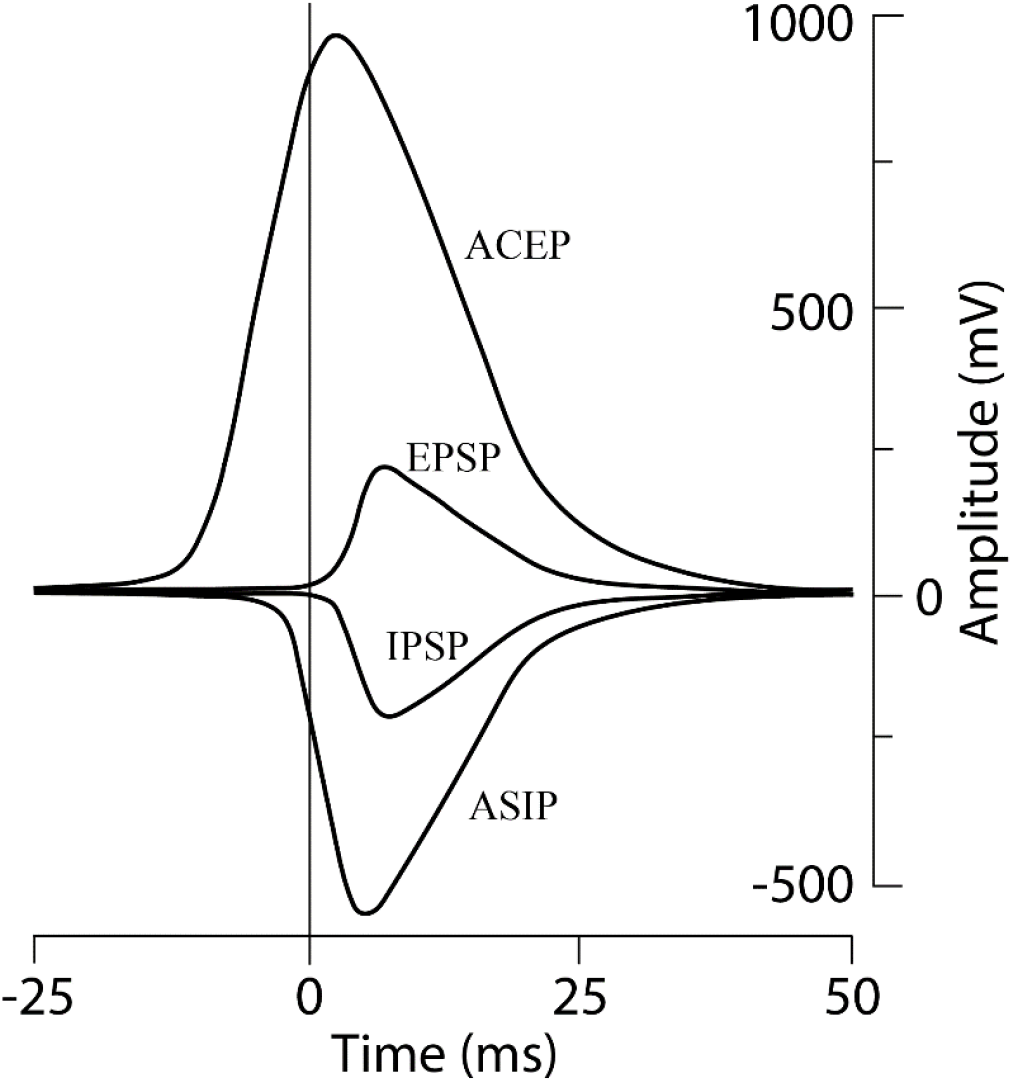
The four types of potentials found in spike-triggered averages of intracellular membrane potentials. The curves were drawn using the average latencies and amplitudes for each type (Table 2 of (Matsumura et al 1996).) Shapes show pure excitatory and inhibitory postsynaptic potentials (EPSPs & IPSPs), with onset after the trigger spike (at T=0), and synchronous excitatory and inhibitory potentials (ASEPs & ASIPs), beginning before the trigger spike. Amplitudes in mV show relative sizes.

Examples of individual STAs showing such synaptic interactions and the sites of the recorded neurons are shown Figs. 3 and 4. Fig. 3 illustrates excitatory synaptic interactions in STAs (right column) between EC and IC neurons whose cortical locations are schematically shown in the left column. The records at top indicate that the EC neuron produced an EPSP in two neighboring IC neurons in the same layer (III). The EPSP in IC1 occurred in isolation, while the EPSP in IC2 was superimposed on an ASEP. The middle records show two examples of ASEP from a layer III EC neuron to two layer V IC neurons. The bottom records show a neuron pair that yielded an EPSP (onset at arrow) superimposed on a slowly rising ASEP.

**Figure 3.**
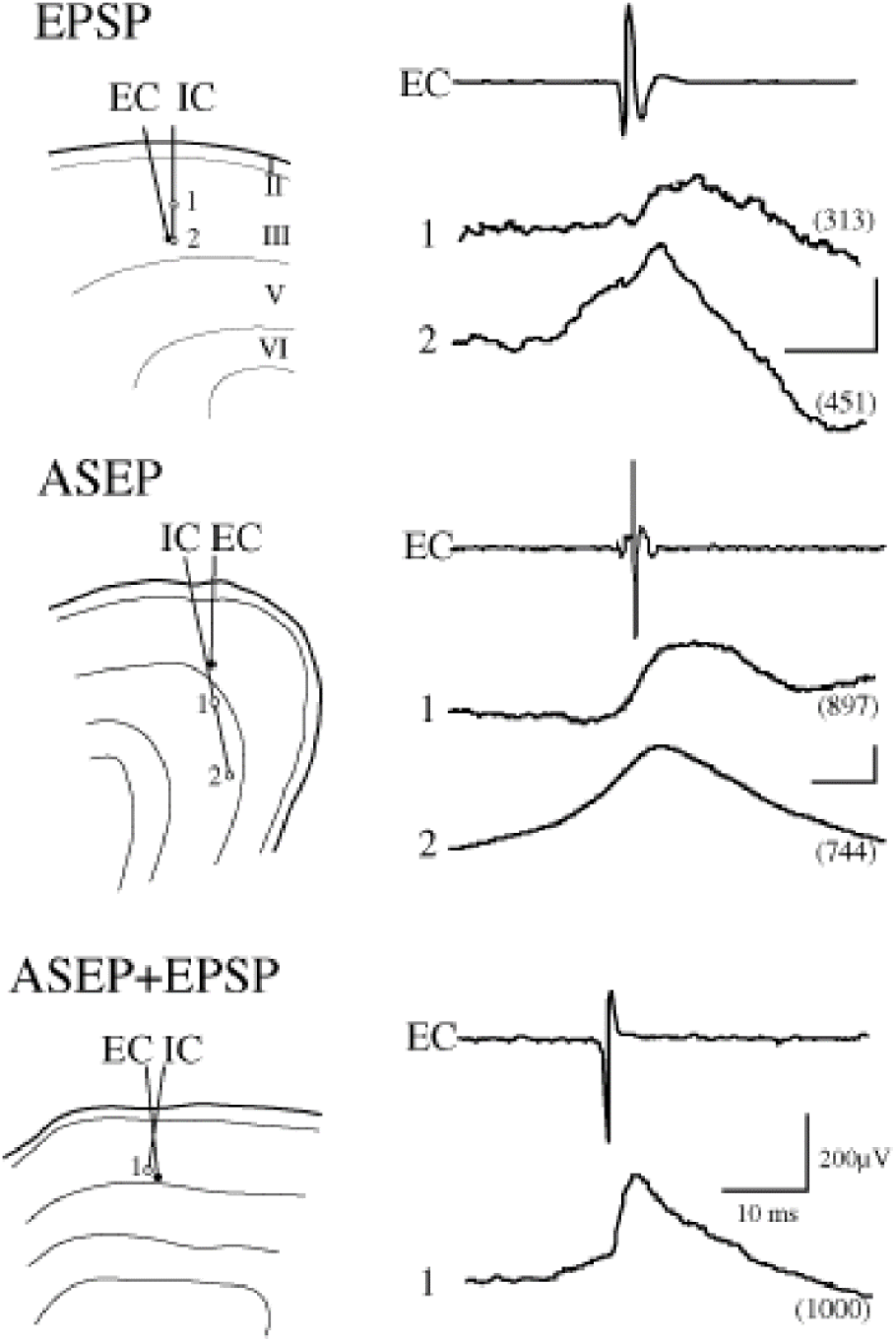
Excitatory synaptic interactions in STAs (right column) and corresponding recording sites (left column). Top: 1. neuron pair (both located in layer III) that yielded the averaged membrane potential featuring EPSP at site 1. 2. ASEP in second IC cell impaled at site 2. Number of sweeps averaged is given in parentheses. Middle: examples of ASEPs from layer III neuron to two layer V neurons. Bottom: a neuron pair that yielded a combination of ASEP and EPSP. Clear deflections are recognized as the onset of ASEP and EPSP (arrows).

**Figure 4.**
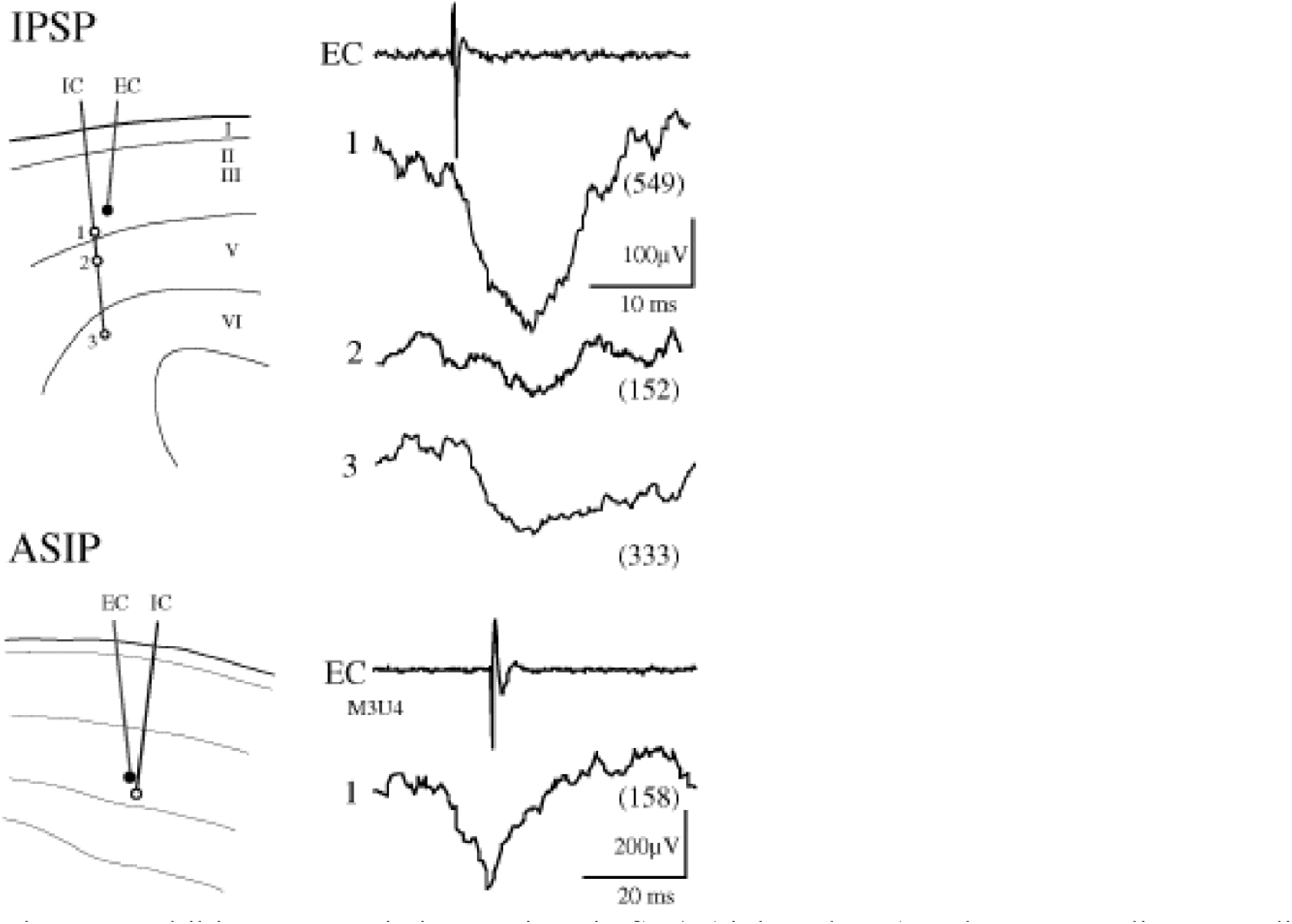
Inhibitory synaptic interactions in STA (right column) and corresponding recording sites (left column). Top: Examples of IPSPs at IC recording sites 1 and 3, shown by downward deflections following the EC spikes. Bottom: Example of ASIP and possible superimposed IPSP.

Figure 4 shows examples of cells with inhibitory synaptic interactions and their cortical locations. Top records show an inhibitory interneuron in layer III that produced IPSPs in two neighboring targets, one in layer III (IC1) and the other in layer VI (IC3). The IC neuron recorded at site 2 did not show any clearly interpretable feature relative to baseline fluctuations. Such ambiguous records were categorized as having no significant feature. The bottom record shows an example of ASIP, whose onset occurred prior to the EC spike. As previously described (Matsumura et al 1996), ASIPs often showed an additional postspike deflection suggesting that the EC cell also produced an IPSP. The sharp negative deflection close to spike onset may reflect an IPSP, but was too ambiguous for confident identification.

As shown in Fig. 4, some STAs were too noisy to allow unambiguous interpretation of features. These were categorized as having no interpretable features, although they might have had some synaptic interactions obscured by synaptic noise, like the example shown here. Since many uninterpretable neuron pairs may actually have had some underlying synaptic interactions, we probably under-estimated the proportion of synaptic connections. In any case, the majority of pairs deemed to have no synaptic interaction had STAs that were completely flat after a large number of sweeps.

### Amplitude of synaptic potentials and direct cell separation

The direct distance between the electrode tips was successfully measured for 220 pairs of EC-IC recordings in the motor and premotor cortex. Of these, 21 showed EPSPs, 135 showed ASEPs, 11 showed IPSPs and 10 showed ASIPs. (The total number exceeds 220 because some recordings contained superimposed features.) The remaining neuron pairs (n=47) were either pairs with no synaptic features or pairs with complex features.

The amplitudes of the synaptic potentials are plotted against the direct distances between the EC and IC neurons in Fig. 5. IPSPs and ASIPs are plotted individually in the same graph (bottom). Amplitudes of both ASEPs (top) and EPSPs (middle) tended to decrease with distance. The ASEP amplitudes between 1.0 to 2.0 mm were smaller than those between 0 to 1.0 mm and between 2.0 to 3.0 mm. For the EPSP group, few neuron pairs could be observed between 1.0 and 2.0 mm, even though many pairs with non-synaptic feature were obtained within this range.

**Figure 5.**
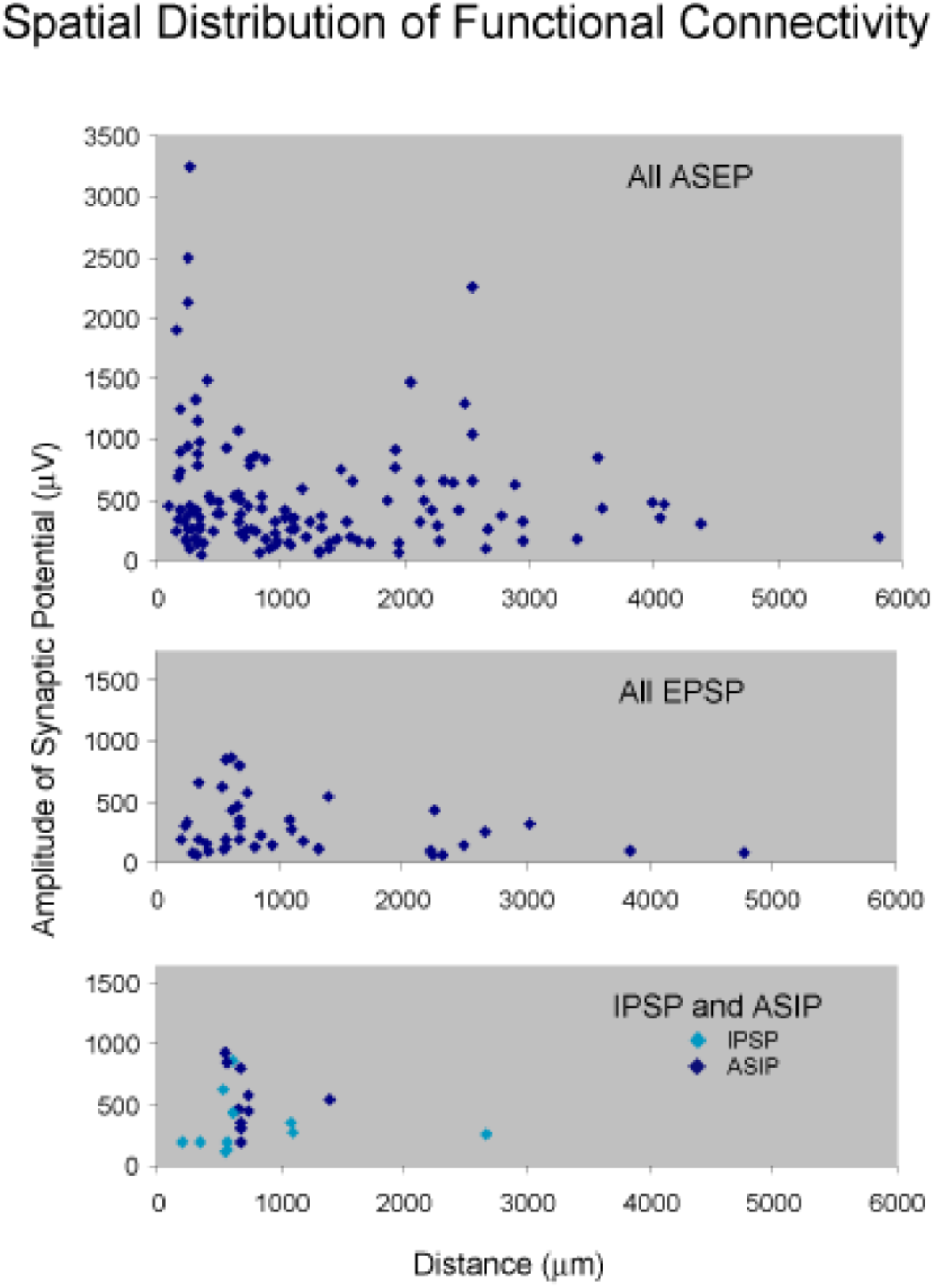
Spatial separation of EC-IC cells and amplitude of synaptic potentials in STAs. Data are grouped as EPSP, ASEP, IPSP, ASIP and all tested neuron pairs that include no feature potentials.

The probability of encountering inhibitory synaptic features decreased when the direct distance between the EC-IC electrode tips exceeded 1.2 mm, though many pairs with no synaptic features were found beyond this distance.

To eliminate possible sampling bias in the recording distances, the normalized probability of each type of synaptic interaction (including no synaptic feature) was plotted against distance (Figure 6). To reduce spatial fluctuations, the probability values were smoothed by calculating the moving average with one neighboring point on each side.

**Figure 6.**
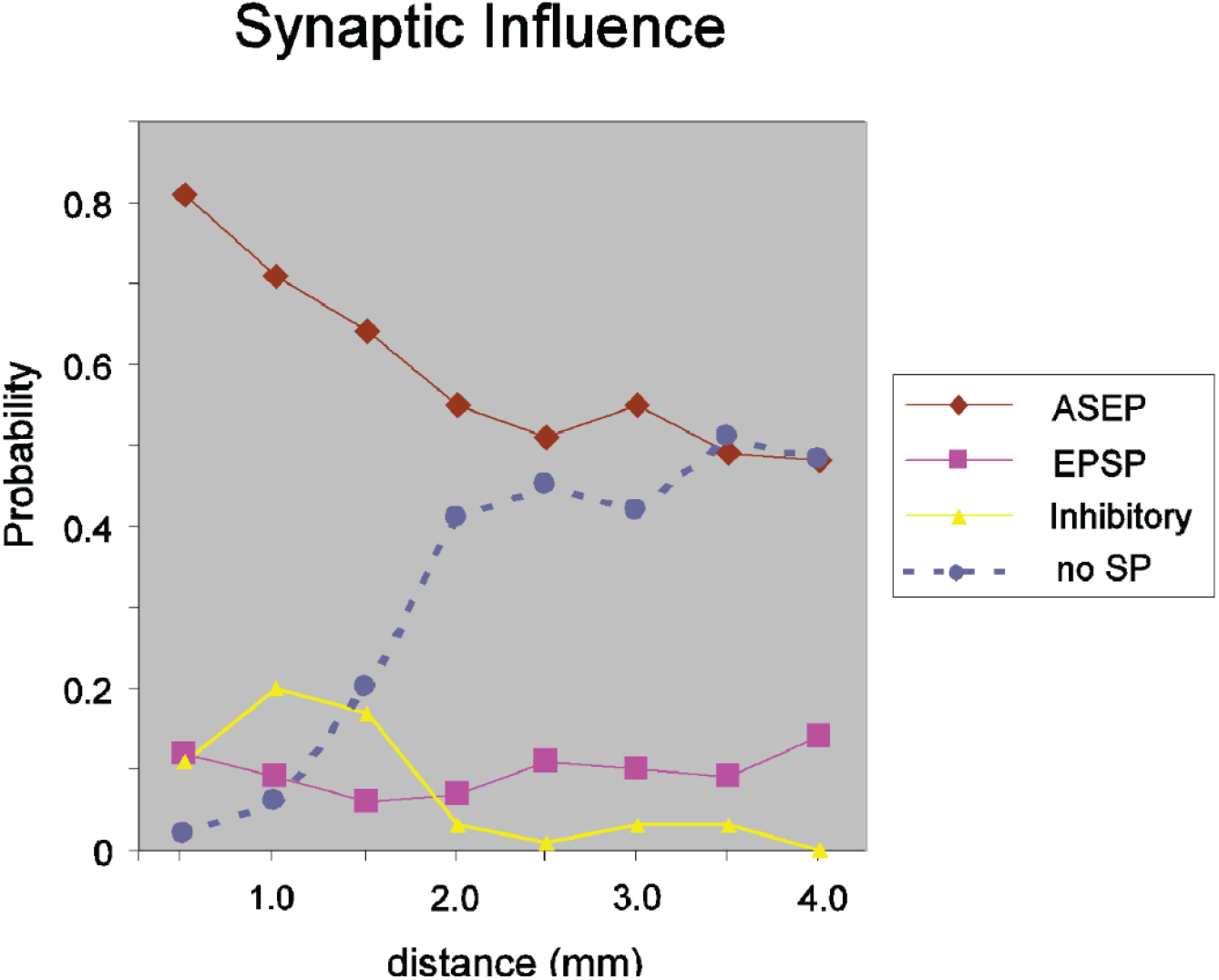
Probabilities of documenting each type of synaptic interaction as a function of cell separation. The probability values were smoothed by averaging neighboring points. When the direct distance was within 0.5 mm, almost all neuron pairs yielded synaptic potential features, primarily ASEPs. Probability of ASEP decayed as EC-IC distance became greater. Compared to the wider distribution of ASEP and EPSP, IPSP and ASIP pairs were largely confined to 1.5 mm. The probability of STAs with no features increased monotonically with distance.

When the direct distance between the EC-IC electrode tips was within 0.5 mm, almost all neuron pairs yielded synaptic potential features. ASEPs were observed for more than 80% of the neuron pairs. As the EC-IC distance increased both the probability and amplitude of ASEPs decreased. In contrast to the wider distribution of ASEP and EPSP, the IPSP and ASIP pairs were more localized, and seldom separated by more than 1.5 mm. The probability of STAs with no features increased with distance. At distances of 3.5 to 4 mm, half of the neuron pairs had no recognizable synaptic features.

### Laminar relationship between the source neuron and target neuron

The laminar distributions for EC-IC neuron pairs that yielded synaptic features were analyzed for 182 neuron pairs that were histologically reconstructed (16 showed EPSPs, 122 ASEPs, 11 IPSPs and 7 ASIPs). The ratio of each feature was comparable with that of 220 neuron pairs described above.

Figure 7 summarizes the proportion of each type of synaptic potential according to the laminar locations of the source and target neurons. ASEP was the most commonly observed synaptic feature in every layer combination. However, for source EC neurons located in layer II-III, this synchronous excitation potential was less frequently observed for target IC neurons in deeper layers.

**Figure 7.**
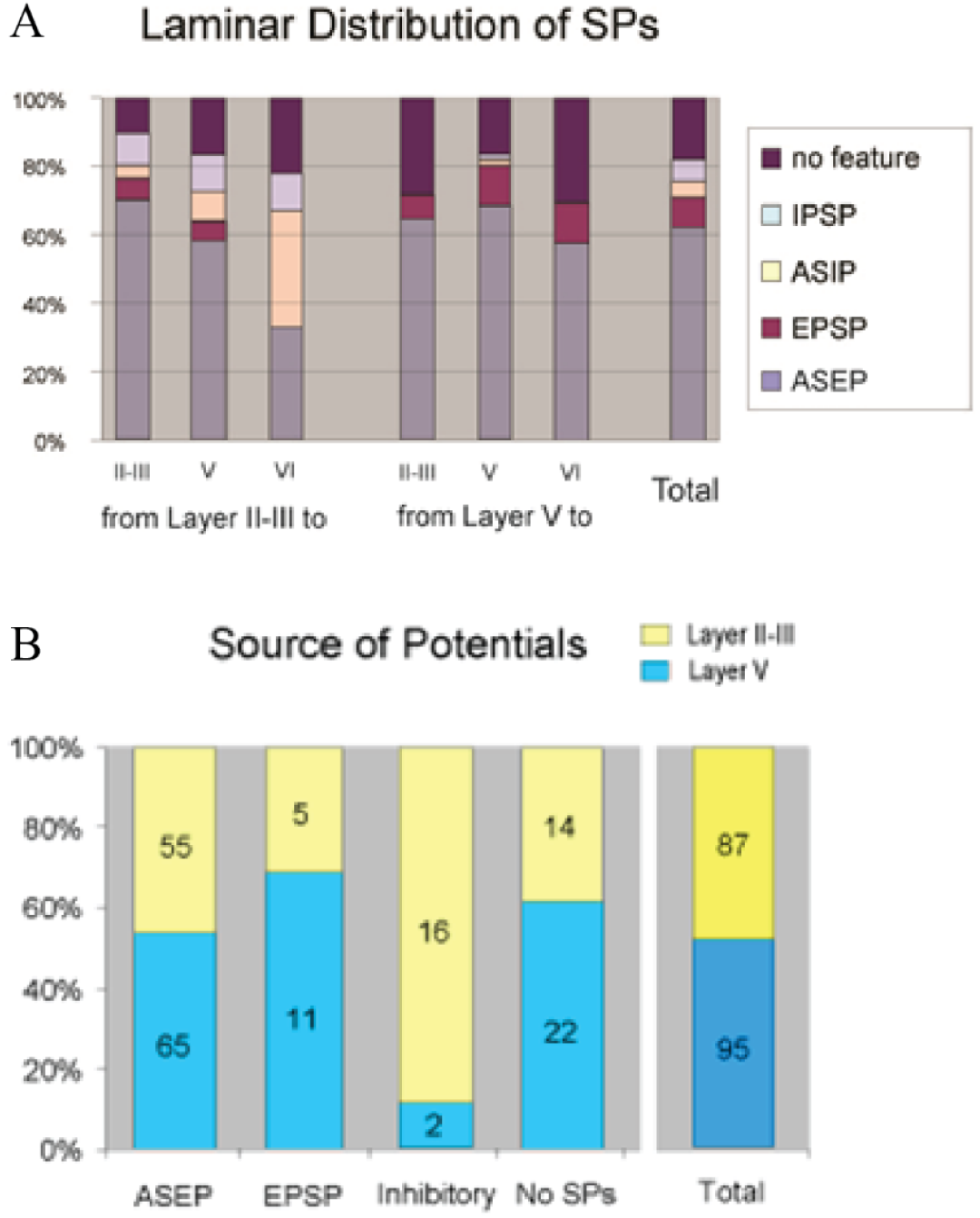
A: laminar distributions for EC-IC neuron pairs that yielded synaptic features. Proportion of each type of synaptic features was summarized, according to the different laminar location of the source and target neurons. In any layer combinations, ASEP was the most commonly observed synaptic feature. However, in cases that the source EC neurons located in layer II-III, these synchronous excitation potentials were less likely observed when the target IC neurons located in deeper layers. Inhibitory features, both IPSPs and ASIPs, were observed seldom among the layer V source EC neurons, suggesting that these inhibitory influences occurred only from superficial to deeper layers. B: IPSP and ASIP apparently dominated in layer II-III source neurons, whereas other synaptic features did not show significant preferences of distribution in between layer II-III and layer V source neurons.

Monosynaptic EPSPs were observed both from upper to lower layers and from lower to upper layers, as well as between neurons in the same layers. In contrast, inhibitory features, both IPSPs and ASIPs, were seldom observed among the layer V source EC neurons, suggesting that these inhibitory influences occurred preferentially from superficial to deeper layers.

A bar graph for EC-IC neuron pairs in the layers II-III and V that yielded each synaptic feature is shown in Fig.7B. IPSP and ASIP apparently dominated in layer II-III source neurons, whereas other synaptic features did not show significant preferential distribution in between layer II-III and layer V source neurons.

### Laminar difference of amplitude vs distance

The amplitudes of the synaptic potentials among EC-IC neuron pairs were further analyzed for laminar differences. Due to the small sample of EPSPs, IPSPs and ASIPs, only ASEPs could be categorized in more detail. Figure 8 shows scatter plots of ASEP amplitudes against direct distances between the EC-IC neuron pairs, separately for the layer II-III source neurons (upper) and for the layer V source neurons (lower). The location of destination IC neurons is color coded. Zero potential values in the graphs indicate neuron pairs with no synaptic features. For pairs from layer II-III to layer V, ASEP amplitude showed large distribution within 1.0 mm. No pairs with larger ASEP amplitude were obtained within a distance between 1.0 to 2.0 mm. However, the amplitude for the pairs between 2.0 to 3.0 mm again showed larger ASEP amplitude similar to those within 1.0 mm. For layer V source neurons, similar discrete distribution of amplitude was seen in a group, projecting to the layer V target neurons.

**Figure 8.**
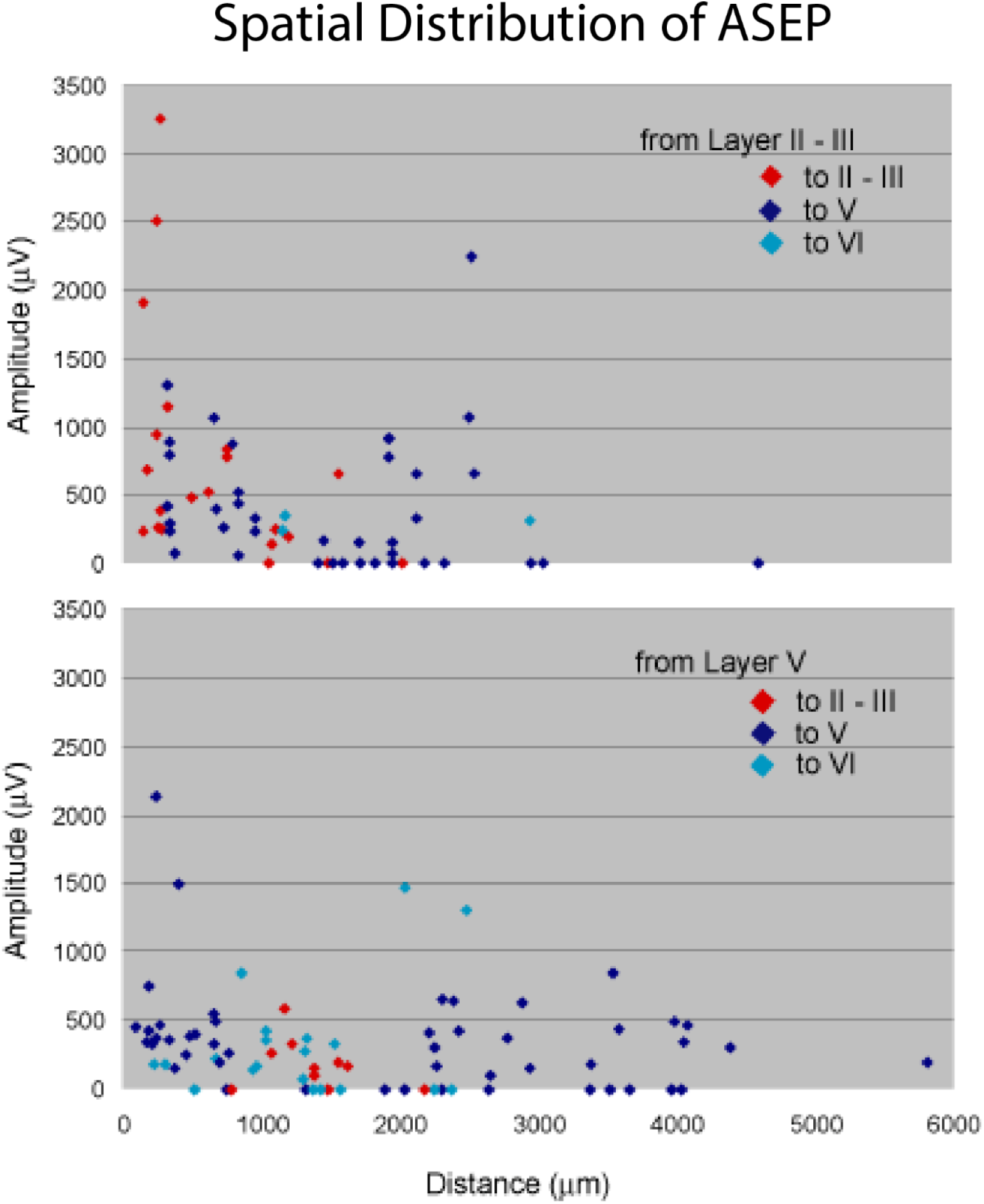
Scatter plots of ASEP amplitudes against direct distances for source neurons in layer II-III (upper) and in layer V (lower), with locations of destination IC neurons color-coded. Zero potential values indicate neuron pairs with no synaptic features.

The distribution of these histologically confirmed neuronal pairs generally resembled the entire population in Fig.6, suggesting that they represented an unbiased sample. The sample of neuronal pairs from layer II-III to layer II-III were largely limited within a distance of 2.0 mm, including non-feature neuron pairs. Within this relatively short range, the amplitude dropped sharply with distance.

For the pairs from layer II-III to layer V, samples were obtained within a distance up to 4.5 mm. Similar to the category of layer II-III to layer II-III, ASEP amplitude showed sharp drop within 1.0 mm. No pairs with larger ASEP amplitude were obtained with separation between 1.0 and 2.0 mm. However, the pairs separated by a distance between 2.0 and 3.0 mm again showed larger ASEP amplitude.

For layer V source neurons, similar discrete distribution of amplitude was seen in a group, projecting to the layer V target neurons. All samples obtained within a distance between 1.0 and 2.0 mm showed no sign of connectivity.

### Horizontal distance vs amplitude of the synaptic features

The periodic changes in amplitude of SPs with separation of the EC-IC neuron pairs, suggest that columnar separation, as well as laminar difference, might influence the amplitude of SPs. We investigated this further by examining the average amplitude of the synaptic potentials for different horizontal separations.

Case numbers of all categorized pairs are shown in Table 1. Unfortunately, one major group, from layer II-III EC to layer II-III IC, lacked samples with wider columnar separation. The only category that we could statistically analyze for the amplitude difference were those of ASEPs from layer II-III to layer V, from layer V to layer II-III, from layer V to layer V, and from layer V to layer VI. As the sample numbers of 2 to 4 columnar separation were not large enough to analyze significance, these data were combined into a single group called *distal* case (Table 2).

**Table 1.**

| Layer Category | ASEP |  |  | EPSP |  |  | ASIP |  |  | IPSP |  |  | No Feature |  |  | Subtotal |  |  |
| --- | --- | --- | --- | --- | --- | --- | --- | --- | --- | --- | --- | --- | --- | --- | --- | --- | --- | --- |
|  | 1 | 2 | >3 | 1 | 2 | >3 | 1 | 2 | >3 | 1 | 2 | >3 | 1 | 2 | >3 | 1 | 2 | >3 |
| II-III to II-III | 21 | 0 | 0 | 2 | 0 | 0 | 1 | 0 | 0 | 3 | 0 | 0 | 3 | 0 | 0 | 28 | 0 | 0 |
| II-III to V | 20 | 12 | 0 | 3 | 0 | 0 | 5 | 0 | 0 | 3 | 3 | 0 | 4 | 5 | 0 | 32 | 19 | 0 |
| II-III to VI | 2 | 1 | 0 | 0 | 0 | 0 | 3 | 0 | 0 | 1 | 0 | 0 | 1 | 1 | 0 | 6 | 2 | 0 |
| V to II-III | 5 | 4 | 0 | 1 | 0 | 0 | 0 | 0 | 0 | 0 | 0 | 0 | 4 | 0 | 0 | 9 | 4 | 0 |
| V to V | 32 | 3 | 6 | 5 | 0 | 2 | 1 | 0 | 0 | 1 | 0 | 0 | 7 | 2 | 1 | 43 | 5 | 8 |
| V to VI | 6 | 9 | 0 | 0 | 0 | 0 | 0 | 0 | 0 | 0 | 0 | 0 | 6 | 2 | 0 | 15 | 11 | 0 |
| **Subtotal** | 86 | 29 | 6 | 11 | 0 | 2 | 10 | 0 | 0 | 8 | 3 | 0 | 25 | 10 | 1 |  |  |  |

*Note: columnar distance of 3 includes three and above*

**Table 2.**

| Amplitudes of ASEP |  |  |  |  |
| --- | --- | --- | --- | --- |
| Columnar Location | II-III to V** | V to II-III | V to V* | V to VI |
| Proximal | 543 | 296 | 457 | 299 |
| Distant | 449 | 153 | 397 | 313 |

Values are mean and SD
\* $p < 0.05$ , \*\* $p < 0.01$

ANOVA showed significant difference of the amplitude of ASEP between the cases of single columnar separation and those of further columnar separation. Further analysis of multiple comparisons showed significant amplitude differences between the *proximal* and *distal* groups in cases from layer II-III to layer V, and from layer V to V. A similar tendency, although not significant, was obtained for the case from layer V to layer II-III, suggesting consistency of reciprocal synchronization between layer II-III and layer V. However, there was no significant difference in a case from layer V to layer VI.

## DISCUSSION

The present study revealed the functional connectivity and spatial distribution of neuronal pairs in the motor cortical areas of monkeys. The strength and polarity of synaptic connections were determined by computing spike-triggered averages of intracellular membrane potentials of the post-synaptic neuron. Before considering functional implications it is important to recognize some sampling biases. The sampled neuron pairs were not uniformly distributed throughout cortical space. Because the two electrodes were advanced through a small opening in the skull, the entry points for both EC and IC electrodes were close, which restricted the separation of the electrode tips at the superficial layers. The electrodes separated further in the depth, allowing sampling of pairs with greater horizontal separation.

The EC and IC electrodes had the usual sampling bias toward recording from large neurons. To the extent that the IC electrode preferably penetrated larger cortical cells, the neuronal sample of IC neurons systematically missed smaller cells and might be different from that of EC neurons. For example, the laminar distribution of the sampled neuron must be carefully examined. Unfortunately, we only obtained one source EC neurons from layer VI, which affected the comparison of the laminar distribution between the source EC neurons and the target IC neurons. The relative ratio of EC and IC recordings between layer II-III and V would give some idea about the sampling bias.

The number of EC neurons recorded from layer II-III was 79, and that from layer V was 69, whereas 41 IC neurons were from layer II-III while 107 were from layer V. This result did not imply immediately that the recording obtained from IC neurons was skewed preferentially to the layer V. As for EC recording, whenever we encountered stable good quality recording, we tried to hold the recording as long as possible while obtaining as many IC recordings from various layers as possible. This caused a sampling bias toward EC neurons in layer II-III. Despite these caveats, the data support several functional conclusions.

Sample bias for the horizontal direction also must be taken account. As for the technical limitation mentioned above, direct distance or columnar separation of layer II-III to layer II-III neuron pairs were no wider than all other laminar combinations. We never had samples with distance of more than 2 mm in this neuron pair category. Nevertheless, even within such a short range of direct distance, a tendency for the amplitude to decay could be recognized, as discussed below.

### Type of synaptic potentials and their laminar distribution

In these experiments, we recorded all four basic synaptic potentials that were previously reported (Matsumura et al 1996). Of all the neuron pairs recorded, approximately 10% yielded EPSPs. Of course, this does not mean that only 10% of the cortical neurons have excitatory axon terminals within the local cortical area. More neurons would send mono-synaptic EPSP to the neighboring neurons. Provided all local neurons send EPSPs, 10% of neurons are expected to be the recipient neurons within a sphere of 4 to 5 mm radius. As more than one third of the cortical neurons are proved to be GABAergic, if the remaining 60% neurons send EPSPs, 17% would be the recipient neurons. The actual connectivity is probably between these two extreme estimations.

Source neurons that send IPSPs altogether with ASIPs, in contrast to EPSPs, showed limited distribution within layer II-III. The recipient neurons, on the other hand, were located widely in all cortical layers. This result confirmed the previous histological studies, in which most of the GABAergic neurons are observed within layer III and send axons to the lower layers as well as layer II-III. Similar estimation attempted for EPSPs would give suggest that GABAergic interneurons send axon terminals to approximately 30% of the neighboring neurons within a sphere of 1.5 mm radius.

Neuron pairs which yielded ASEP had the widest laminar distribution. Since there are no source-target relations for the synchronous activity between neuron pairs, it could be inferred that layer VI neurons have synchronized potentials with many neurons in layer II-III to layer V neurons, as their relationship is reciprocal. ASEP was the most common synaptic drive in the motor cortex, consistent with observations in other cortical areas. Many neurons that showed ASEP had other synaptic features such as EPSP, IPSP and even ASIP. It is noteworthy that more than 50% of the neuron pairs within a sphere of 4.5 mm radius shared common synaptic drive through the superficial and deep cortical layers.

### Separation of excitatory and inhibitory synchronous connections

Separation of the neuron pairs was related to the amplitude of the synaptic potentials differently depending on the type of the SPs and their laminar location. ASEP had the highest synaptic connectivity among all other features, and more than 80% of the neuron pairs received commonly driven excitatory inputs if the direct distance was within 0.5 mm with highest amplitude of the peak STAs, as shown in Figs 6 and 8.

ASEP dominated among various synaptic influences within a separation less than 1.0 mm. But ASEP almost suddenly disappeared between a distance range of 1.0 mm and 2.0 mm, and reappeared over 2.0 mm, as shown in Fig.8. This implies that there might be a functional separation or independence in directly adjacent columns. Functional similarity seems to re-appear for separation over 2.9 mm. Evidence of similar spatially periodic interactions has been reported in cross-correlation studies in visual cortex (Ts’o et al 1986)

Several models of the intrinsic connectivity of neurons in visual cortex have been proposed (Douglas & Martin 1991, Douglas & Martin 2004, Toyama et al 1981) One can question whether circuitry of visual cortex, designed to analyze sensory input, would be organized similarly to motor cortex circuits designed to generate motor output. Our results are consistent with the proposed visual cortex models, but the details need to be further explored, with greater sampling of neuronal pairs.

### Functional roles of cortical micro-circuit

The contrast between the wide distribution of cells with excitatory synaptic connections (ASEP and EPSP) and the more localized distribution of cells with inhibitory links (IPSP and ASIP) suggests that the latter may be involved in sculpting functional columns. Some of the inhibitory connections are intercolumnar, serving to separate activity in adjacent columns and mediating surround inhibition. However, these inhibitory connections could also be intracolumnar and thereby involved in neural calculations within a column. This issue can be further investigated by documenting the response properties of the connected neurons.

## ACKNOWLEDGEMENTS

This work was supported by NIH grants NS12542 from NINDS and RR00166. We thank Mr. Larry Shupe for technical support.

